# scPheno: A deep generative model to integrate scRNA-seq with disease phenotypes and its application on prediction of COVID-19 pneumonia and severe assessment

**DOI:** 10.1101/2022.06.20.496916

**Authors:** Feng Zeng, Xuwen Kong, Fan Yang, Ting Chen, Jiahuai Han

**Affiliations:** Department of Automation, Xiamen University, Xiamen, Fujian 361102, China; Department of Neuroscience, School of Medicine, Xiamen University, Xiamen, Fujian 361102, China; Department of Neurosurgery, The First Affiliated Hospital of Xiamen University, Xiamen University, Xiamen, Fujian, China; State Key Laboratory of Cellular Stress Biology, School of Life Sciences, Xiamen University, Xiamen 361005, China; National Institute for Data Science in Health and Medicine, Xiamen University, Xiamen 361005, China; Institute for Artificial Intelligence, Department of Computer Science and Technology, Tsinghua University, Beijing 100084, China; Tsinghua-Fuzhou Institute of Digital Technology, Beijing National Research Center for Information Science and Technology, Tsinghua University, Beijing 100084, China

## Abstract

Cell-to-cell variability is orchestrated by transcriptional variations participating in different biological processes. However, the dissection of transcriptional variability in specific biological process at single-cell level remains unavailable. Here, we present a deep generative model scPheno to integrate scRNA-seq with disease phenotypes to unravel the invisible phenotype-related transcriptional variations. We applied scPheno on COVID-19 blood scRNA-seq to separate transcriptional variations in regulating COVID-19 host immunity and transcriptional variations in maintaining cell-type identity. *In silico*, we found CLU^+^IFI27^+^S100A9^+^ monocyte as the efficient cellular marker for the prediction of COVID-19 diagnosis. Inspiringly, using only 4 genes upregulated in CLU^+^IFI27^+^S100A9^+^ monocytes can predict the COVID-19 diagnosis of individuals from different country with an accuracy up to 81.3%. We also found C1^+^CD163^+^ monocyte and 8 C1^+^CD163^+^ monocyte-upregulated genes as the efficient biomarkers for the prediction of severity assessment. Overall, scPheno is an effective method in dissecting the transcriptional basis of phenotype variations at single-cell level.

## Main Text

Coronavirus disease 2019 (COVID-19) pneumonia, caused by the infection of a novel severe acute respiratory syndrome coronavirus 2 (SARS-CoV-2), is a unprecedented global crisis. On one side, it is reported that 40.5% of individuals are with asymptomatic infection^1^, which poses a threat to public health. On the other side, a considerable number of the mild/moderate patients will deteriorate to severe or even critical status rapidly^2^, which puts a high pressure on scarce health resources. Developing the effective strategy of diagnosis and early warning of severity is one of the major challenges of the COVID-19 pandemic, which is valuable for clinical management and decision making.

The diagnosis and severity assessment of COVID-19 primarily use chest X-ray imaging and chest computed tomography (CT) imaging, which provide the visualization of lung injury mediated by the invasion of SARS-CoV-2 virus^3^. However, when the disordered structures of the SARS-CoV-2 virus-infected lung are large enough for visualizing and detecting, the deteriorating patients may miss opportunities for early intervention. Recent advances in single-cell RNA sequencing (scRNA-seq) promote the inspection of how immune system acts against the invading SARS-CoV-2 virus. The large-scale single-cell studies such as COMBAT^4^ (COvid-19 Multi-omics Blood Atlas Consortium) and SC4^5^ (Single Cell Consortium for COVID-19 in China) reveal that SARS-CoV-2 virus changes the cellular composition of immune system systematically^6^. First, COVID-19 is marked with a remarkable decline in the numbers of circulating macrophages, dendritic cells, natural killer (NK) cells, T cells, and B cells. Second, host immunity displays several abnormal phenomena in correlation with COVID-19 severity^2^, such as emergency myelopoiesis^7^ triggered by acute inflammatory response against SARS-CoV-2 that gives rise to massive expansion of immature neutrophils^8^ and dysfunctional monocytes^4^. Third, the adaptive immune compartment demonstrates the atypical phenotypes too. For example, it is controversial whether the PD-1-expressing T cells elevated in COVID-19 patients are exhausted or functional^9,10^. These findings suggest that the characteristics of host immune abnormality extracted from blood scRNA-seq could serve as new predictive biomarkers in addition to chest X-ray images and chest CT images. However, it remains underestimated the value of blood scRNA-seq at the development of new diagnostic and therapeutic strategies.

Dissection of the variations on cell state and transcriptome between different conditions is critical for the identification of disease-associated biomarkers. To date, the methods of identifying the disease-associated biomarkers mainly rely on the changes in cell population abundances across pathological conditions^11–13^. In essence, this kind of computational approaches assumes that the effect of disease on cell populations is discrete, deterministic, and instantaneous. However, the selective pressure exerted by disease shapes cell state and transcriptome consecutively. For example, it is reported that the myeloid lineage in COVID-19 exhibits significant plasticity^7^. Therefore, the identification of the COVID-19-associated biomarkers should consider the continuous variations on gene expression and cell state apart from the discrete changes in cell population abundances. Unfortunately, it is still lacking that a method associates the continuous gene expression variations with discrete disease phenotypes at single-cell level.

In this paper, we present a method based on deep generative probabilistic model^14^ for separating gene expression variations associated with disease phenotypes from gene expression variations associated with cell fates, estimating the specificity of disease phenotype for each cell, and predicting the disease phenotypes of individuals based on the ensemble of the association strengths of immune cells with disease phenotypes. Our method can be used to diagnose various disease phenotypes such as health, influenza, sepsis, and COVID-19, and can be used for the stratification of COVID-19 patients, evaluating the illness severity from the asymptomatic status to critical status. Moreover, with the use of deep generative probabilistic modeling, our method enables the integration of the scRNA-seq datasets of large-scale COVID-19 cohorts to provide a comprehensive immune cell atlas for finding the efficient cellular and molecular markers for the prediction of COVID-19 diagnosis and severity assessment.

### Overview of the cell type-phenotype association analysis

Overall, our method is made up of the information extraction and disease prediction components (Fig. 1a). First, in the information extraction component, we define the association strengths between immune system and diseases by the joint distribution of cell states and disease phenotypes of immune cells, and design a deep generative probabilistic model called scPheno to model the joint distribution explicitly (Fig. 1b). Both disease phenotype and cell fate decision impact the transcriptional states of cells. Thus, it is possible to predict the cell-type identity and disease phenotype from gene expression simultaneously. We assume that gene expression variations modulating the variability between immune cells can be decomposed into two mutually independent parts, of which the first part accounts for the determination of cell state and the second part accounts for the cell-to-cell variability on disease phenotype. scPheno models the joint distribution of cell states and disease phenotypes by involving two deep probabilistic models to describe the distribution of gene expression variations associated with cell states and the distribution of gene expression variations associated with disease phenotypes, respectively (Fig. 1c). On one side, scPheno can predict the cell states (or cell types) of immune cells to gauge the cell type composition of immune system. On the other side, scPheno can predict the disease phenotypes of immunes cells to quantify the strength of each immune cell expressing gene programs specific to a disease. We conducted a series of experiments and demonstrated that scPheno is accurate at predicting the cell-types and phenotypes of cells (Supplementary text, Supplementary Figs. 1 to 7). Second, in the disease prediction component, our method uses the joint distribution of cell states and disease phenotypes of immune cells as the predictive features and uses support vector machine (SVM) for the classification of disease phenotypes of patients.

**Fig. 1.**
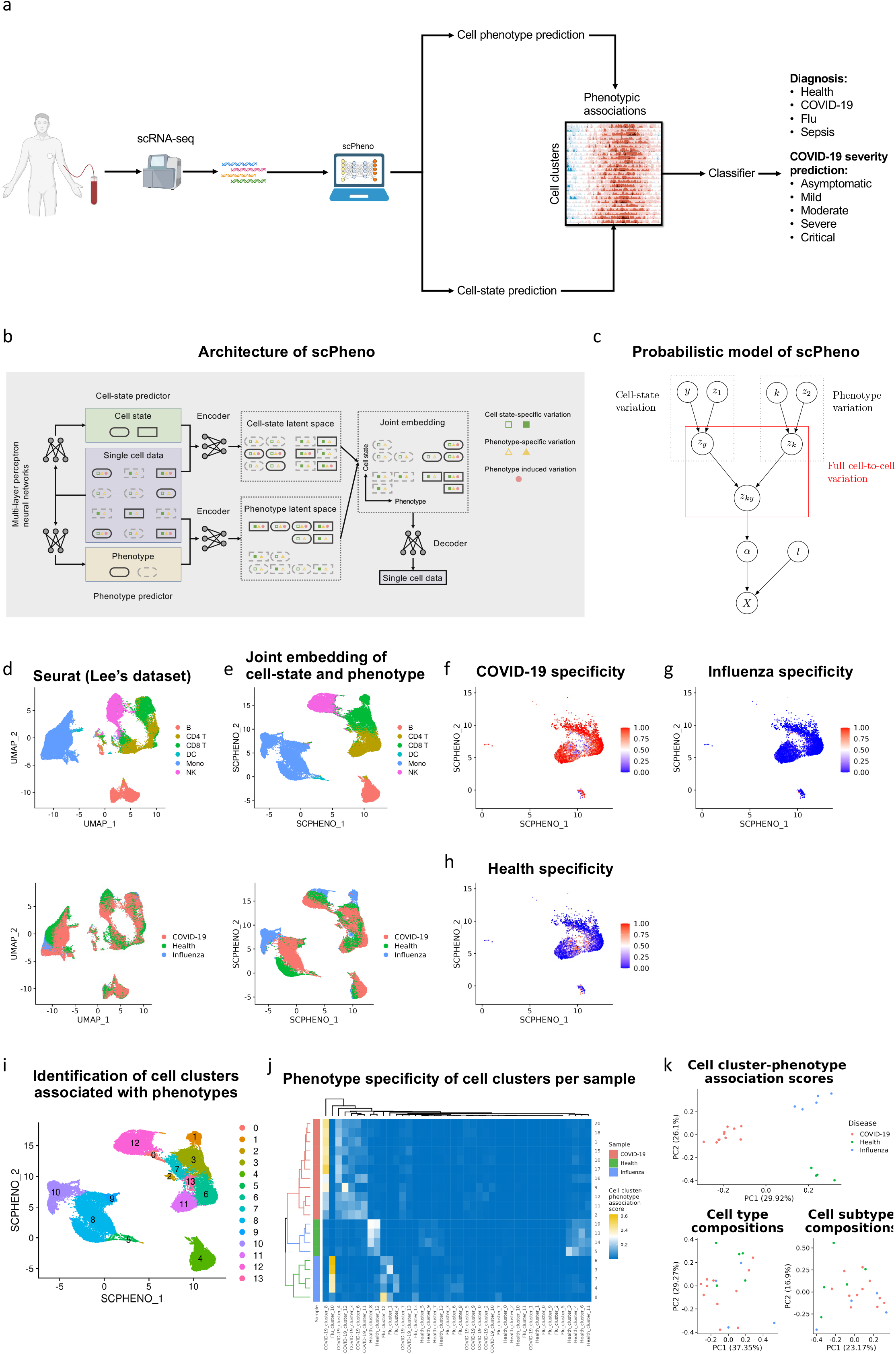
COVID-19 diagnosis and severity prediction based on the covariation of cell states and disease phenotypes of immune cells. **a**, The workflow of COVID-19 diagnosis and severity prediction. **b**, The architecutre of scPheno for the decomposition of transcriptional variations associated with cell states and disease phenotypes. **c**, The probabilistic model of scPheno for describing the full cell-to-cell variability. **d**, The UMAP visualization of Lee‘s dataset. **e**, The visualization of full cell-to-cell variability in the joint embedding space. **f**, COVID-19 specificity scores of CD4^+^ T cells in COVID-19 patients. **g**, Influenza specificity scores of CD4^+^ T cells in COVID-19 patients. **h**, Sepsis specificity scores of CD4^+^ T cells in COVID-19 patients. **i**, The determination of cell clusters. **j**, The aggregation of samples based on the cell state-disease phenotype associations. **k**, The PCA plots based on the cell state-disease phenotype associations, the cell-type composition, and the cell-subtype composition.

### scPheno maximizes the dissection of cell-to-cell variability with respect to the variations on cell type and phenotype

As a proof-of-concept experiment, we applied our method on Lee’s dataset^15^. Lee’s dataset contains gene expression profiles of 63,509 immune cells in peripheral blood samples collected from 4 healthy individuals, 5 influenza patients, and 11 COVID-19 patients. Firstly, we used Seurat^16^ to process the single-cell gene expression profiles and visualized the distribution of cell states and disease phenotypes with the uniform manifold approximation and projection (UMAP) method^17^ (Fig. 1d). Unfortunately, it is hard to identify the variability of immune cells on disease phenotypes. Next, we analyzed the single-cell gene expression profiles with scPheno. scPheno obtained several advances. On one side, scPheno generated a better population structure of distinct immune cell states in the cell state latent space (Supplementary Fig. 10a). On the other side, in the cell phenotype latent space, it is more convenient to identify the differences of immune cells associated with disease phenotypes. For example, it is clearly displayed that monocyte cells have three distinct subpopulations associated with healthy status, influenza disease, and COVID-19 disease, respectively (Supplementary Fig. 10b). Furthermore, through the combination of the above two latent spaces, scPheno revealed the complete cell-to-cell variability on both cell state and disease phenotype simultaneously (Fig. 1e). Even within the seemingly homogeneous cells, scPheno can leverage the phenotype specificity score to uncover the transcriptional differences among the cells. For example, scPheno reveals the subtle heterogeneity of CD4^+^ T cells in COVID-19 patients. CD4^+^ T cells with high COVID-19 specificity scores indicate that they express the COVID-19-specific genes (Fig. 1f). The rest of CD4^+^ T cells with low COVID-19 specificity scores are more like those seen in healthy individuals (Figs. 1g and h). These results validate our assumption that transcriptional variations between cells are arisen from various sources, correlate with different biological processes, and can be separated from each other.

Owing to the decomposition of transcriptional variations associated with cell states and transcriptional variations associated with disease phenotypes, scPheno improves the inspection resolution of cell-to-cell variability. Based on the joint embedding of cell states and disease phenotypes, cell clusters that are more specific to disease phenotypes can be identified to improve the stratification of patients (Fig. 1i). For example, we identified new cell clusters for Lee’s dataset, and precisely separated individuals with different disease phenotypes according to the association strengths of cell clusters with disease phenotypes (Fig. 1j). The principal component analysis (PCA) on the heatmap of cell cluster-phenotype association confirmed the results (Fig. 1k). We also conducted PCA by using the cell-type composition and the cell-subtype composition as comparisons. Consequently, it is unable to separate COVID-19 patients from healthy individuals and influenza patients according to the differences of either the cell-type composition or the cell-subtype composition (Fig. 1k).

### A large-scale blood immune cell atlas of COVID-19

We used scPheno to integrate the COMBAT dataset^4^, Haniffa’s dataset^18^, and Lee’s dataset^15^, and established a large-scale immune cell atlas in blood from multiple pathological conditions and from different locations (Fig. 2a). Because scPheno models gene usages instead of modeling UMI counts, it can effectively remove batch effects in scRNA-seq datasets (Supplementary text, Supplementary Figs. 8 and 9). The integrated atlas comprises a total of 1,378,873 immune cells constituting a detailed continuum of transcriptional variations in immune system challenged by various kinds of viruses and bacteria (Fig. 2b). It contains 46 healthy individuals, 17 influenza patients, 37 sepsis patients, 13 asymptomatic COVID-19 patients, 56 mild COVID-19 patients, 43 moderate COVID-19 patients, 70 severe COVID-19 patients, and 42 critical COVID-19 patients in several regions and countries (Fig. 2a). The integrated immune cell atlas reveals that T cell, B cell, NK cell, and monocyte compartments exhibit the phenotype-specific patterns (Supplementary Fig. 11).

**Fig. 2.**
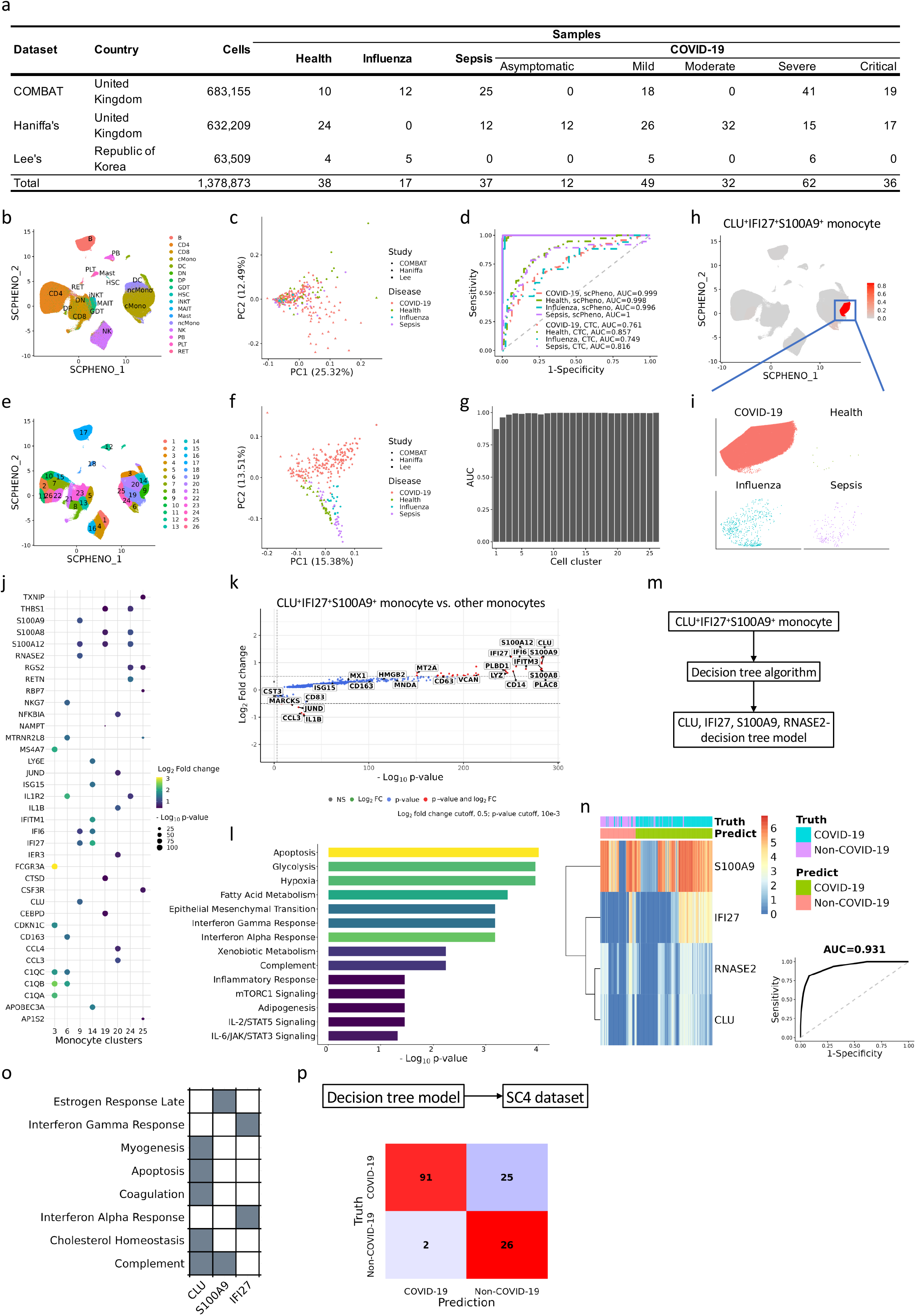
Establishment of COVID-19 immune cell atlas, evaluation of COVID-19 diagnosis prediction, and the search for predictive biomarkers. **a**, The statistics of three scRNA-seq datasets being integrated. **b**, The visualization of the integrated immune cells in the joint embedding space. **c**, The PCA visualization of the samples based on the cell-type composition analysis. **d**, The ROC curves and AUC scores of scPheno versus the cell-type composition analysis. **e**, The determination of cell clusters. **f**, The PCA visualization of the samples based on the scPheno analysis. **g**, Selection of the predictive subset of cell clusters with the recursive feature elimination (RFE) algorithm. **h**, CLU^+^IFI27^+^S100A9^+^ monocyte is effective at predicting COVID-19 diagnosis. **i**, Distribution of CLU^+^IFI27^+^S100A9^+^ monocyte cells across disease phenotypes. **j**, Top 6 differentially expressed genes in every monocyte clusters. **k**, Volcano plot of differentially expressed genes between CLU^+^IFI27^+^S100A9^+^ monocyte cells and other monocyte cells. CLU^+^IFI27^+^S100A9^+^ monocyte cells upregulate the reported COVID-19-associated genes including VCAN, HMGB2, MT2A, MX1, PLBD1, and PLAC8. **l**, Pathway enrichment analysis of CLU^+^IFI27^+^S100A9^+^ monocyte-specific genes with the MSigDB Hallmark Pathway database. **m**, Build a decision tree model from the CLU^+^IFI27^+^S100A9^+^ monocyte-specific genes. **n**, Heatmap visualization and ROC curve of the prediction of the samples in COVID-19 immune cell atlas with the decision tree model. **o**, MSigDB Hallmark pathways that CLU, IFI27, S100A9, and RNASE2 are involved in. **p**, The confusion matrix when applying the decision tree model on the SC4 dataset.

We first predicted COVID-19 and other phenotypes based on the immune cell-type composition, which is a commonly used prediction strategy in single-cell studies. To resolve the conflicts in the cell-type nomenclature between datasets, cells of the Haniffa’s dataset and Lee’s dataset were aligned onto the COMBAT dataset and assigned the identities according to the cell-type nomenclature used in COMBAT with scPheno. It has been enormously documented that the changes of various immune cell types correlate with COVID-19. But the PCA analysis showed that COVID-19 samples were not separated from samples of other phenotypes based on the composition of immune cell types (Fig. 2c). The receiver operator characteristics (ROC) curve was obtained with the leave-one-out crossvalidation (LOOCV) strategy. The area under the ROC curve (AUC) of disease prediction based on the immune cell composition was 0.796 ± 0.050 on average (Fig. 2d). The separation of COVID-19, influenza, and sepsis is difficult, because mild COVID-19 displays the flu-like manifestations, e.g. cough and fatigue, and severe COVID-19 and sepsis have the similar fatal symptoms. Next, we determined cell clusters in the integrated immune cell atlas (Fig. 2e), and performed the PCA analysis based on the cell cluster (type)-phenotype association scores. It is noticeable that the samples of different phenotypes were well separated (Fig. 2f). scPheno improves the prediction of disease phenotypes significantly. The average AUC of disease prediction was increased up to 0.998±0.002 (Fig. 2d).

### CLU^+^IFI27^+^S100A9^+^ monocyte and a subset of 4 genes can predict COVID-19 diagnosis accurately

Though the above results are inspiring, it is clinically impracticable to predict COVID-19 based on the blood single-cell transcriptomics due to the limitations of time and cost. Thus, we asked whether there exists a subset of cells or a subset of genes with the similar prediction power as that of using the blood single-cell transcriptomics. To answer this question, we used the recursive feature elimination (RFE) algorithm to assess the prediction powers of the possible combinations of cell clusters (Fig. 2g). It is supervising that COVID-19 could be predicted with the accuracy of 0.872 when only one cell cluster was used. The most powerful predictive cluster was the cluster 9 (Figs. 2h and i). Compared with other monocyte clusters, the cluster 9 specifically expresses the apoptotic gene CLU, the interferon-stimulated genes IFI6 and IFI27, and the alarmins S100A9 and S100A12 (Fig. 2j). The cluster 9 is termed as CLU^+^IFI27^+^S100A9^+^ monocyte throughout the paper. The functional enrichment analysis revealed that the upregulated genes in CLU^+^IFI27^+^S100A9^+^ monocytes are enriched in apoptosis, interferon alpha response, interferon gamma response, and complement reaction (Figs. 2k and l).

For every individual, we summarized over the cells to create the pseudo-bulk RNA-seq data. Then, the decision tree algorithm was run to search for a subset of genes upregulated in CLU^+^IFI27^+^S100A9^+^ monocytes, which could be as the efficient gene markers for the prediction of COVID-19 diagnosis (Fig. 2m). This yielded a decision tree model with four genes, which are CLU, IFI27, S100A9, and RNASE2 (Fig. 2n, Supplementary Fig. 12). It is noticed that CLU and S100A9 participate in complement reaction (Fig. 2o). We applied the four-gene decision tree model onto the SC4 dataset, which is comprised of individuals all recruited in China. It is encouraging that the simple decision tree model with only four genes can obtain an accuracy of 0.819 at the prediction of COVID-19 diagnosis for the unseen samples (Fig. 2p).

### C1^+^CD163^+^ monocyte and a subset of 8 genes can predict COVID-19 severity assessment accurately

Next, we evaluated the prediction of COVID-19 severity assessment. For simplicity, we used the term mild/moderate to represent the patients with the asymptomatic, mild, and moderate phenotypes, and used the term severe/critical to represent the patients with the severe and critical phenotypes. We compared the compositions of immune cells in the mild/moderate patients and the severe/critical patients (Fig. 3a). Overall, the abundance of classical monocyte cells was elevated in the severe/critical patients, and the abundances of CD4^+^ T cells and CD8^+^ T cells were elevated in the mild/moderate patients (Supplementary Fig. 13). However, the PCA analysis revealed that the differences on the cell-type compositions between mild/moderate and severe/critical were not effective in predicting COVID-19 severity assessment (Figs. 3b and f). But the transcriptional states of all immune cell types changed in correlation with COVID-19 severity (Fig. 3c). scPheno combined the cell-type information and the transcriptional information and succeeded at separating the mild/moderate patients and the severe/critical patients (Figs. 3d and f). The Bayes factor analysis revealed that the changes on the transcriptional states of monocytes were most correlated with COVID-19 severity (Fig. 3e). Like the above analysis, we found that using only the cluster 6 can obtain an AUC score of 0.903 at predicting the COVID-19 severity assessment (Fig. 3g, Supplementary Fig. 14). The cluster 6 differentially expressed several complement genes including C1QB and C1QC and the antiinflammatory gene CD163 (Fig. 2j). Thus, the cluster 6 was termed as C1^+^CD163^+^ monocyte throughout the paper. Pathway enrichment analysis revealed that the upregulated genes in C1^+^CD163^+^ monocytes were enriched in complement reaction and interferon gamma response (Supplementary Fig. 15). We also performed the aforementioned decision tree algorithm, and found a predictive decision tree model built from a subset of eight C1^+^CD163^+^ monocyte-upregulated genes (Fig. 3h). The eight genes were ACTB, MYOF, NCF1, DYNLT1, TNFSF10, GIMAP7, LDHA, and MIF. We applied the 8-gene decision tree model on the SC4 dataset and obtained an prediction accuracy of 0.707 (Fig. 3i).

**Fig. 3.**
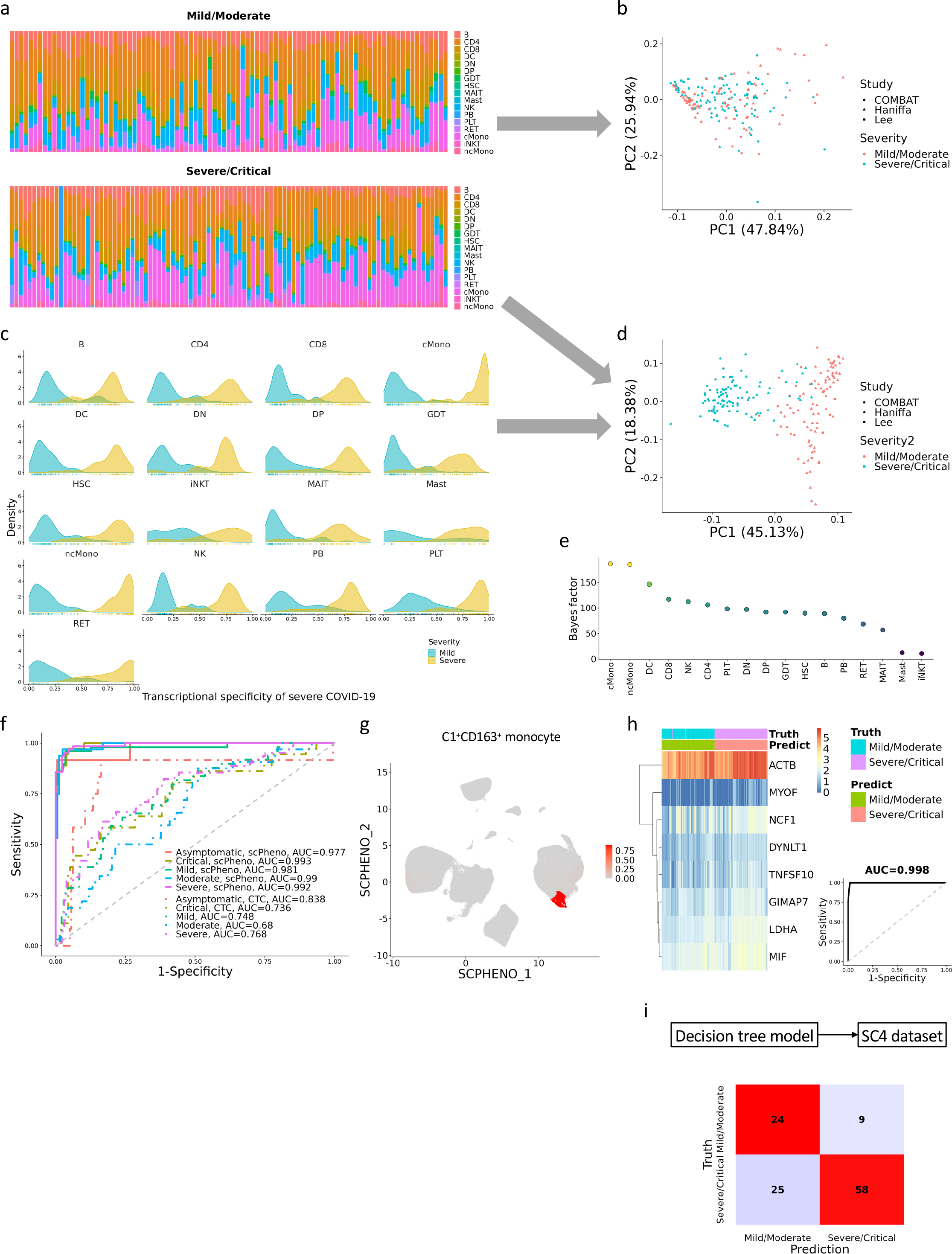
Evaluation of COVID-19 severity assessment prediction and the search for the predictive biomarkers. **a**, Barplots of the cell-type compositions of the mild/moderate patients and the severe/critical patients. **b**, The PCA visualization of the mild/moderate patients and the severe/critical patients based on the cell-type composition analysis. **c**, The strengths of immune cell types expressing the COVID-19-specific gene expression programs in the mild/moderate patients and the severe/critical patients. **d**, The PCA visualization of the mild/moderate patients and the severe/critical patients based on the combination of the cell-type information and the phenotype-specificity information. **e**, COVID-19 specificity scores of samples derived from the ensemble of cell phenotype scores. **f**, The ROC curves and AUC scores of scPheno versus the cell-type composition analysis. **g**, C1^+^CD163^+^ monocyte is effective at predicting COVID-19 severity assessment. **h**, Heatmap visualization and ROC curve of the prediction of the samples in COVID-19 immune cell atlas with the decision tree model built from the C1^+^CD163^+^ monocyte-specific genes. **i**, The confusion matrix when applying the decision tree model on the SC4 dataset.

## Discussion

It is a key question in life sciences how isogenic cells act in different ways under the same pressure. The cell-to-cell variability is not accessible by bulk RNA-seq. Recently, with the advent of scRNA-seq, a unprecedented number of cell heterogeneity has been reported. The scRNA-seq studies of COVID-19 also revealed that the infection of SARS-CoV-2 virus contributes to substantial changes in immune cells with different outcomes. Meantime, the scRNA-seq studies of COVID-19 provided the elaborate medical records of patients’ phenotypes. This gives a great opportunity for deciphering the transcriptional basis of the phenotypic variability of seemingly homogeneous cells. Mapping from transcriptome to phenotype is a challenge problem. No computational tools currently exist for this kind of task.

The transcriptional variation of a cell is complicated, which is a consequence of the coordinated dynamics of numerous biological processes. In this paper, we conjecture that it is possible to decompose the transcriptional variations associated with different biological processes. It is noticed that each biological process can shape the transcriptional state and induce variations into the transcriptome. Conversely, it is possible to predict the biological process from the variations on the transcriptome. Hence, we formulated the decomposition of transcriptional variations as the classification problems. For instance, scPheno is to classify the cell types and to classify the phenotypes simultaneously. With the classification objective and deep generative modeling, it is possible to capture the transcriptional variations associated with different biological processes, that is, *p*(*x*|*k*) where *x* denotes transcriptional state and *k* biological process.

We propose a machine-learning framework for the implementation of the above idea, and provide a computational tool scPheno to extract transcriptional variations associated with disease phenotypes from single-cell transcriptomics for the dissection of cell heterogeneity at high resolution. We applied scPheno on the blood scRNA-seq of COVID-19, and found two subpopulations of monocytes that were effective in predicting COVID-19 diagnosis and severity assessment. We also found two subsets of predictive genes. The result is inspiring that it may be feasible to develop the clinically testing device with a few predictive genes mined from blood single-cell transcriptomics for rapid, accurate, and non-invasive COVID-19 diagnosis and severity assessment in the future.

Our method can be deployed on personal computer equipped with GPU device to process scRNA-seq datasets comprised of millions of cells within hours (Supplementary text, Supplementary Figs. 3, 5, and 7). One limitation of our method is that multiple phenotypes are not supported. The limitation can be resolved by representing individual’s phenotypes with one-hot encoding. In the future, we will adopt the one-hot encoding to support the assessment of the effects of multiple phenotypes, e.g., age, gender, race, and comorbidity on cell heterogeneity. Although application demonstrations are about COVID-19, the proposed method can be used in the single-cell study of any diseases and cancers for the integration of patient information with scRNA-seq.

## Methods

### scPheno

Gene expression profile observed in a cell is resulted from the mixture of various kinds of transcriptional variations, reflecting the effects of diverse biological processes on the state of the cell. We hypothesize that biological processes are distinguishable and identifiable, and thereafter the biological process-specific transcriptional variations can be decomposed out of the scRNA-seq data. To this end, we designed a deep probabilistic model called scPheno (short for **S**ingle **C**ell **PHENO**type) to formulate the generation of the single-cell transcriptional observations resulted from the collection of variations on cell states within individual biological processes. For the simplicity in the rest of the paper, only the biological processes of cell-type determination and phenotype change are considered in our method. But our model can be easily extended to involve more complicated biological processes.

Following our hypothesis, the generation procedure of single-cell gene expression is a hierarchical process. First, conceptually, given a cell, the hidden states of the cell associated with the cell-type determination and phenotype change are determined. Second, the hidden states of two biological processes are combined to form the full cell state. Third, gene expression profile of the cell is generated according to the full cells state.

In mathematical terms we define the variables *y* and *k* to denote cell type and cell phenotype, respectively. Cell type is the identity of the cell. It can be the grained cell nomenclatures such as T cell and B cell, and can also be the fine cell nomenclatures such as subtypes and clusters. The resolution of cell type depends on the input annotations given by users. Cell phenotype means the conditions under which the cells are collected, e.g., illness situations and experimental perturbations. In scPheno, both cell type *y* and cell phenotype *k* are modeled as the categorical variables. The variables *z_y_* and *z_k_* are introduced into represent the hidden states associated with the cell-type determination and phenotype change, respectively. These two hidden state variables are generated by the following supervised probabilistic principal component analysis (SPPCA) models^21^. Specifically, for the hidden state variable *z_y_*, the model is

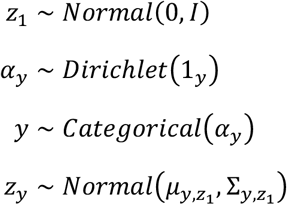

where, 1_*y*_ = [1,⋯,1] is the parameter of the prior distribution over *α_y_*, ||1_*y*_|| = *n_y_* is the number of cell types. The prior distribution implies that cells of different cell types occur with equal probability. The vector *μ*_*y,z*_1__ = [*μ*_*y,z*_1_,1_,⋯,*μ*_*y,z*_1_,h_] and the diagonal matrix 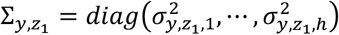 represent the mean and covariance of *z_y_*, where *h* is the dimension of *z_y_*. In scPheno, *μ*_*y,z*_1__ and ∑_*y,z*_1__ are the nonlinear functions of *y* and *z*_1_, implemented by a multiple layer perceptron (MLP) neural network. Likewise, for the hidden state variable *z_k_*, the model is

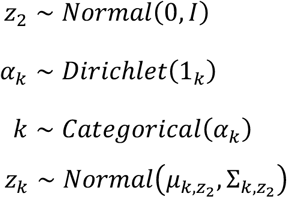

where, 1_*k*_ = [1,⋯,1] is the parameter of the prior distribution over *α_k_*, ||1_*k*_|| = *n_k_* is the number of phenotypes of interest. The vector *μ*_*k,z*_2__, = [*μ*_*y,z*_2_,1_,⋯,*μ*_*k,z*_2_,*h*_] and the diagonal matrix 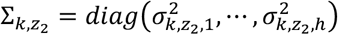 represent the mean and covariance of *z_k_*, and are approximated by a MLP neural network with *k* and *z_2_* as the input. The full cell state is defined as *z_ky_* = *z_k_*⊙*z_y_*, where the operator ⊙ represents the concatenation of two vectors.

Next, it is to model the generation of the single-cell observation *X* given the full cell state *z_ky_*. Due to shallow sequencing depth, the variance of single-cell gene expression measurements is greater than the expectation, which is known as over-dispersion statistically. It remains challenging to produce the accurate estimates of gene expression values from the over-dispersed counts of unique molecular identifiers (UMIs). Though many statistical techniques such as the zero-inflation models and negative binomial models have been widely used to cope with the great variability in UMI counts, these models assume that genes are independent from each other. As a matter of fact, genes interact with each other and participate in various gene expression programs which are necessary for the maintenance of cellular functionality. The frequency of genes used in gene expression programs defines gene usage in a cell, which characterizes the correlation structure in gene expression. Gene usage is intrinsic to cell and is not affected by technical biases, including batch effects and inflated zeros (that is dropouts). Therefore, scPheno is to model gene usage instead of UMI counts. Let *θ* = *θ*_1_, ⋯, *θ_n_g__*] denote gene usage, 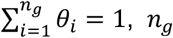, *n_g_* is the number of genes. We also introduce a factor to describe the effect of library size denoted by the variable *l.* Hence, the generation of single-cell gene expression *X* is as follows,

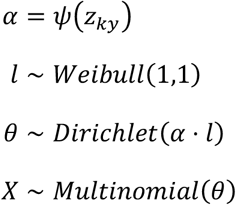

where, *ψ* is implemented with a MLP neural network. Taken together, the joint probability of the single-cell observation and unknown variables is

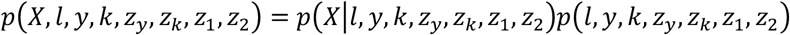

where, *p*(*l,y,k,z_y_,z_k_,z*_1_, *z*_2_) represents the prior probability over unknown variables, and *p*(*X*|*l, y, k, z_y_, z_k_,z*_1_, *z*_2_) represents the probability of observing *X* when the cell type *y* and cell phenotype *k* are given.

Variational inference is effective for the learning of the complicated probabilistic models. To use variational inference, we define the variational distribution as *q*(*l,y,k,z_y_,z_k_,z*_1_,*z*_2_|*X*) to approximate the true distribution *p*(*l,y,k,z_y_,z_k_*,*z*_1_,*z*_2_). According to the mean-field theory, the variational distribution can be decomposed into

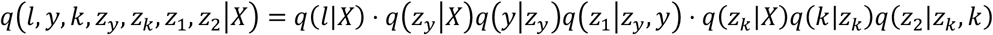

where,

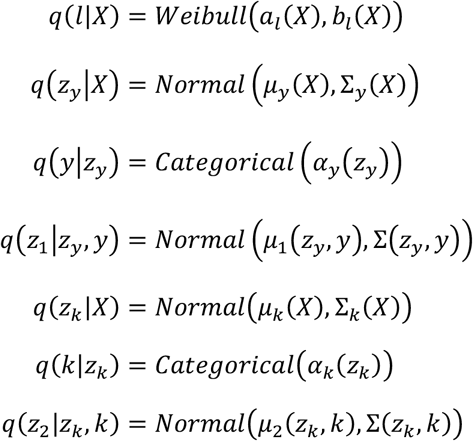

where, the parameters of the above decomposed variational distribution models are approximated by the MLP neural networks. For the simplicity in the rest of the paper, all the unknown variables are collectively represented by the latent variable *Z* = (*l, y, k, z_y_, z_k_, z*_1_, *z*_2_).

The loss function for learning parameters is made up of three components. First, it is anticipated that the variational distribution *q*(*Z*|*X*) approximates *p*(*Z*) as closely as possible. The divergence between *q*(*Z*|*X*) and *p*(*Z*) is measured by using the Kullback– Leibler divergence *KL*(*q*(*Z*|*X*)||*p*(*Z*)). Second, the latent variable *Z* drawn from the variational distribution *q*(*Z*|*X*) can optimally explain the generation of the single-cell observation *X* with the proposed deep probabilistic model *p*(*X*|*Z*). Therefore, it is anticipated that the expectation of data likelihood *E*_*q*(*Z*|*X*)_[*p*(*X*|*Z*)] should be maximized. The combination of the Kullback–Leibler divergence and expected data likelihood is known as the evidence lower bound (ELBO). Third, it is anticipated that the variational models *q*(*y*|*z_y_*) and *q*(*k*|*z_k_*) can predict cell type and phenotype accurately. Cross-entropy is used to define the losses of cell-type classification and phenotype classification, that is, 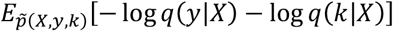, where *q*(*y*|*X*) = ∫ *q*(*y*|*z_y_*)*q*(*z_y_*|*X*)*dz_y_* and *q*(*k*|*X*) = ∫ *q*(*k*|*z_k_*)*q*(*z_k_*|*X*)*dz_k_*. Taken together, the total loss function is

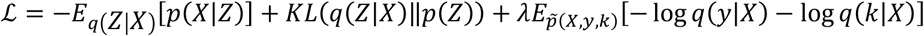

where, 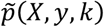 is the empirical distribution of single-cell observations with cell type annotations and phenotype annotations. The hyperparameter *λ* impacts the effectiveness of the cell-type classification and phenotype classification. In order to obtain the good enough classifiers, the hyperparameter *λ* should be set as the value that makes the negative ELBO and cross-entropy loss with the same order of magnitude.

In implementation, all MLP neural networks are three-layered neural networks with the same number of hidden neurons. The optimal parameters of the MLP neural networks used in both generative models and variational models are estimated with the ADAM optimization algorithm.

### Low-dimensional representation of single cells and the prediction of both cell types and phenotypes

With the variational distribution models, scPheno can be used for multiple tasks of scRNA-seq data analysis. First, scPheno provides various kinds of low-dimensional representations of single cells for the visualization of cell-to-cell variability from different aspects. The variational distribution model *q*(*z_y_*|*X*) is to visualize the variation of cells in the cell-type latent space. The variational distribution model *q*(*z_k_*|*X*) is to visualize the variation of cells in the phenotype latent space. The concatenation of *z_y_* and *z_k_* yields the cell type-phenotype latent variable *z_ky_* = *z_k_*⊙*z_y_* that is to visualize the full cell-to-cell variability.

Second, the variational distribution model *q*(*y*|*X*) = ∫ *q*(*y*|*z_y_*)*q*(*z_y_*|*X*)*dz_y_* can be used to predict the cell-type *y* given gene expression signature of a cell *X.* Likewise, the variational distribution model *q*(*k*|*X*) = ∫ *q*(*k*|*z_k_*)*q*(*z_k_*|*X*)*dz_k_* can be used to predict the phenotype of a given cell *X.*

### Disease prediction and severity prediction of individuals

For the prediction of disease status and illness severity, we introduce the joint distribution *p*(*y,k*) to represent the predictive feature of an individual, which is the covariation of cell types *y* and phenotypic conditions *k.* The joint distribution *p*(*y,k*) is approximated through the summation of the cell types and phenotypes of all cells in an individual, that is,

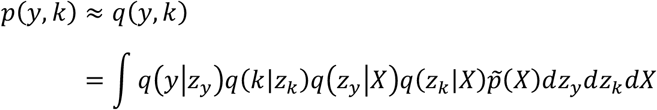

where 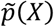 is the empirical distribution of single-cell gene expression signatures. SVM is used for disease prediction and severity prediction, where the input features are the joint distributions *p*(*y,k*).

For comparison, we also predict disease status and illness severity based on either the abundances of cell types in an individual or the abundances of cell subtypes in an individual. The prediction machine is SVM.

### Performance evaluation

There are some difficulties at evaluation. First, sample sizes of the COVID-19 scRNA-seq datasets are small. Second, the number of COVID-19 samples and the number of samples with other phenotypes are not balanced. To circumvent the difficulties, we adopted the leave-one-out cross-validation (LOOCV) strategy. Specifically, at each iteration, it used one sample as the validation data, trained the evaluating method with the rest of samples, and applied the well-trained method to predict the validating sample. The process was repeated until all samples have been validated. The prediction results were collected to compute the area under the receiver operator characteristic curve (AUC) for the evaluation of COVID-19 diagnosis and severity prediction. The receiver operator characteristic (ROC) curve is a systematic assessment of a classification method, illustrating how the true positive rate and false positive rate change along with different thresholding values. AUC is an overall score of the performance of a classification method.

### Recursive feature elimination, SVM, Bayes factor, and decision tree

The analyses of recursive feature elimination, SVM, and Bayes factor were performed with the public Python package and R package. The recursive feature elimination analysis used the RFE function in the Python package Scikit-learn^22^ with the default parameters. The SVM analyses used the SVC function in the Python package Scikit-learn with the default parameters. The Bayes factor analysis use the regressionBF function in the R package BayesFactor with the default parameters. The decision tree analysis used the function rpart in the R package rpart with the parameter minsplit=5.

### COVID-19 scRNA-seq datasets and data processing

The COVID-19 scRNA-seq datasets used in this paper were fetched from the public data repositories. The COMBAT dataset was downloaded from the Zenodo website (https://doi.org/10.5281/zenodo.6120249). The Haniffa’s dataset was retrieved from the COVID-19 Cell Atlas website (https://www.covid19cellatlas.org). The Lee’s dataset and the SC4 dataset were accessible at the NCBI GEO database with the accession numbers GSE149689 and GSE158055.

scRNA-seq datasets were processed following the instructions of the R package Seurat. The UMI counts were normalized and scaled with the functions NormalizeData and ScaleData. The highly variable genes (HVGs) were determined with the function FindVariableFeatures. The function RunUMAP was used to compute the UMAP visualization.

## Code availability

TBD.

## Acknowledgements

F.Z. was supported by National Natural Science Foundation of China (61503314) and Natural Science Foundation of Fujian Province, China (2019J01041).

## Author contributions

F.Z. and J.H. conceived the concept and supervised the project. F.Z. designed the model. F.Z. and X.K. wrote the code. F.Z., X.K. performed the experiments. F.Z., X.K., F.Y., T.C., and J.H. analyzed the experiment results.

---

**Supplementary Fig. 1.**
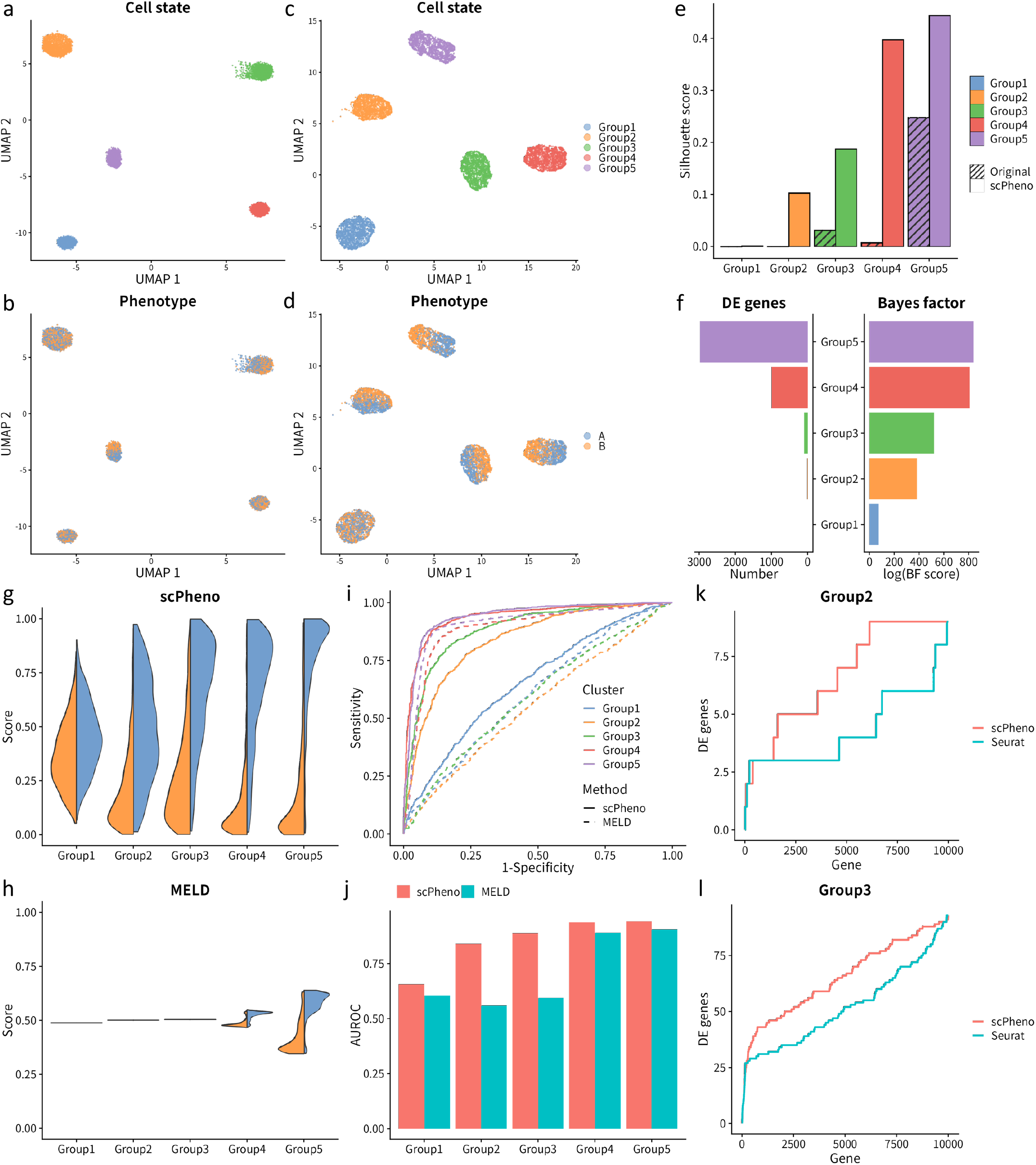
Evaluation of phenotype heterogeneity analysis with the simulation scRNA-seq dataset. **a**, The visualization of cell states of the simulated dataset when the UMAP algorithm was applied on the PCA embedding yielded by Seurat which only considered cell types. **b**, The visualization of phenotypes of the simulated dataset when the UMAP algorithm was applied on the PCA embedding yielded by Seurat which only considered cell types. **c**, The visualization of cell states of the simulated dataset when the UMAP algorithm was applied on the cell type-phenotype embedding yielded by scPheno. **d**, The visualization of phenotypes of the simulated dataset when the UMAP algorithm was applied on the cell type-phenotype embedding yielded by scPheno. **e**, Ranking of immune cell types based on the Bayes factor analysis on the transcriptional specificity relating to COVID-19 severity. **f**, The numbers of phenotype associated differentially expressed (DE) genes in cell clusters, and the prioritization of cell clusters based on the phenotype specificity indicated by the Bayes factor analysis. **g**, The distribution of the phenotype likelihood scores of cells yielded by scPheno. **h**, The distribution of the phenotype likelihood scores of cells yielded by MELD. **i**, The ROC curves of predicting the phenotypes of cells. **j**, The AUC scores of predicting the phenotypes of cells. **k**, The frequency of the phenotype associated DE genes detected at the ranking positions for Group2. **l**, The frequency of the phenotype associated DE genes detected at the ranking positions for Group3.

**Supplementary Fig. 2.**
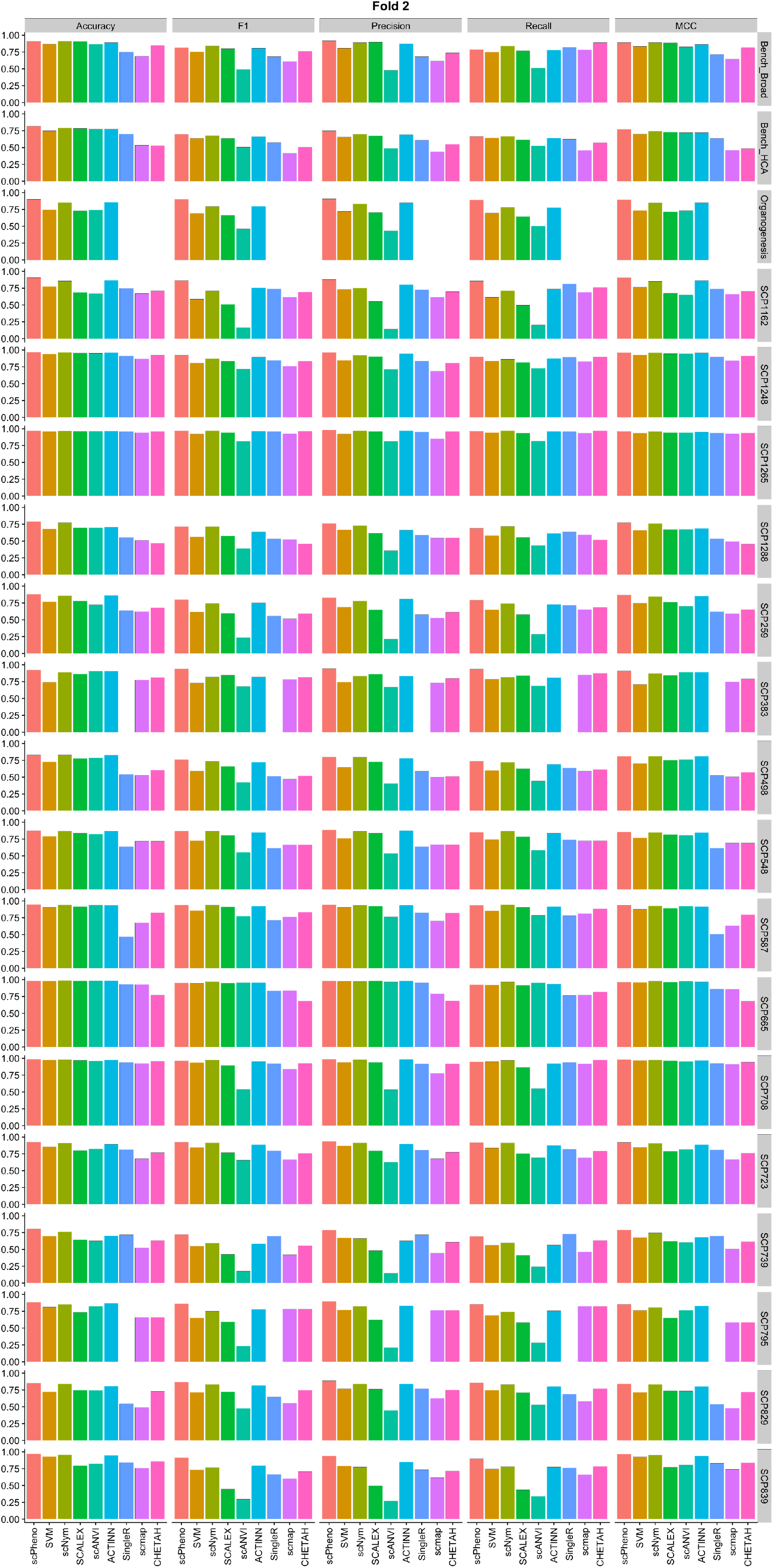
Evaluation of the cell-type prediction, two-fold. For each benchmark dataset, 50 percent of the dataset were used for validation and the rest for training.

**Supplementary Fig. 3.**
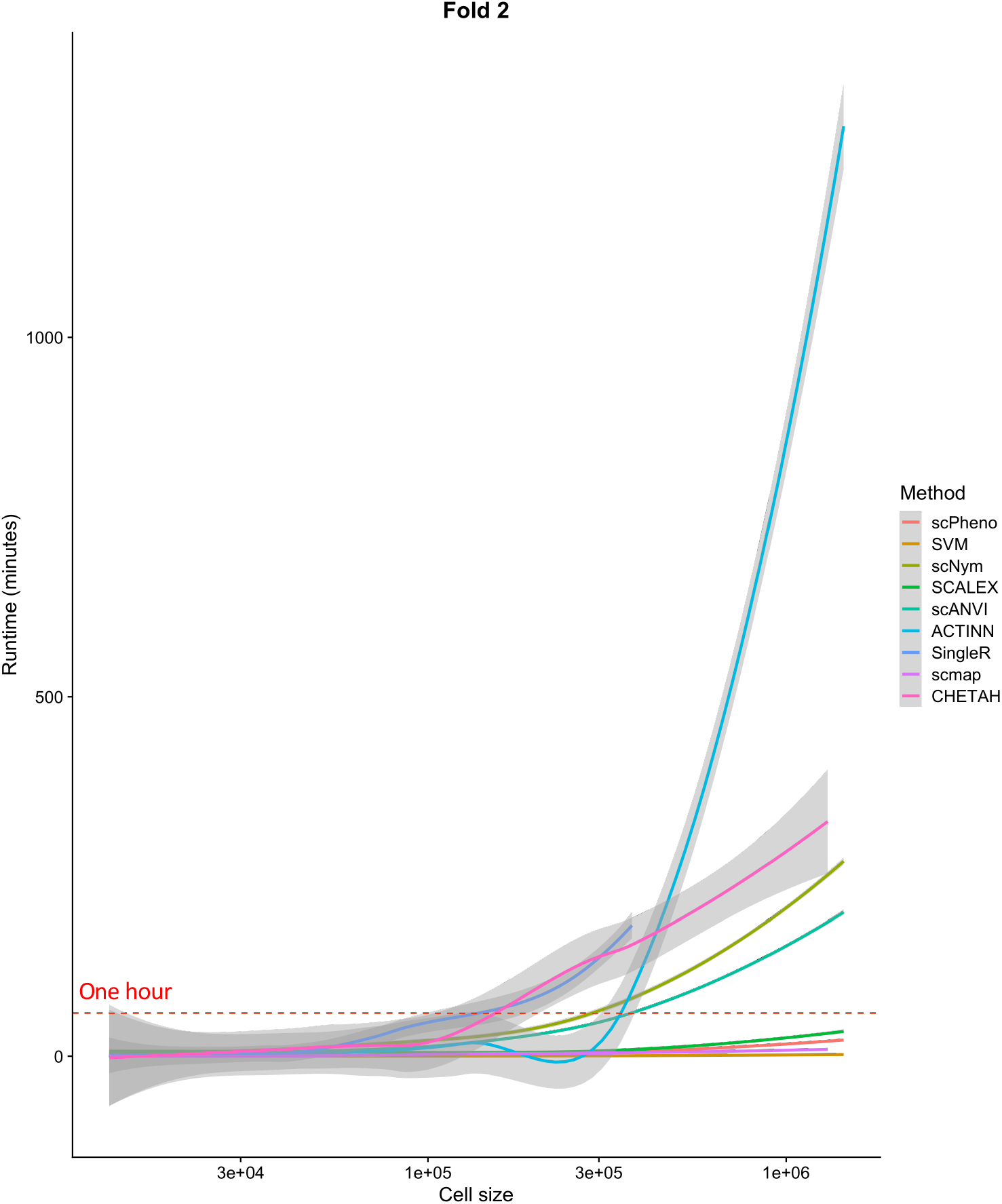
Runtime of the two-fold evaluation experiment including the training time and the validation time. The red dashed line represents one hour.

**Supplementary Fig. 4.**
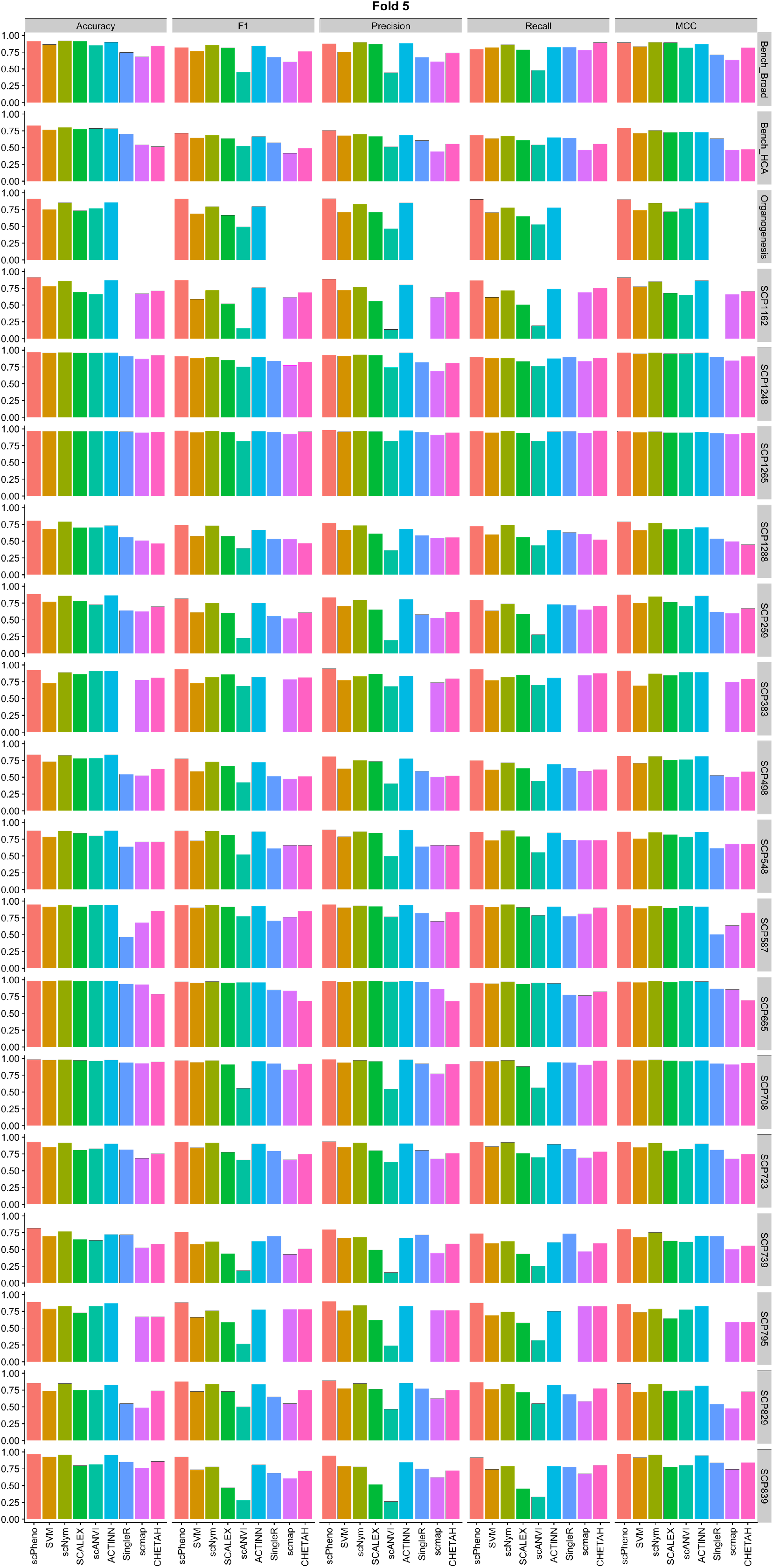
Evaluation of the cell-type prediction, five-fold. For each benchmark dataset, 80 percent of the dataset were used for validation and the rest for training.

**Supplementary Fig. 5.**
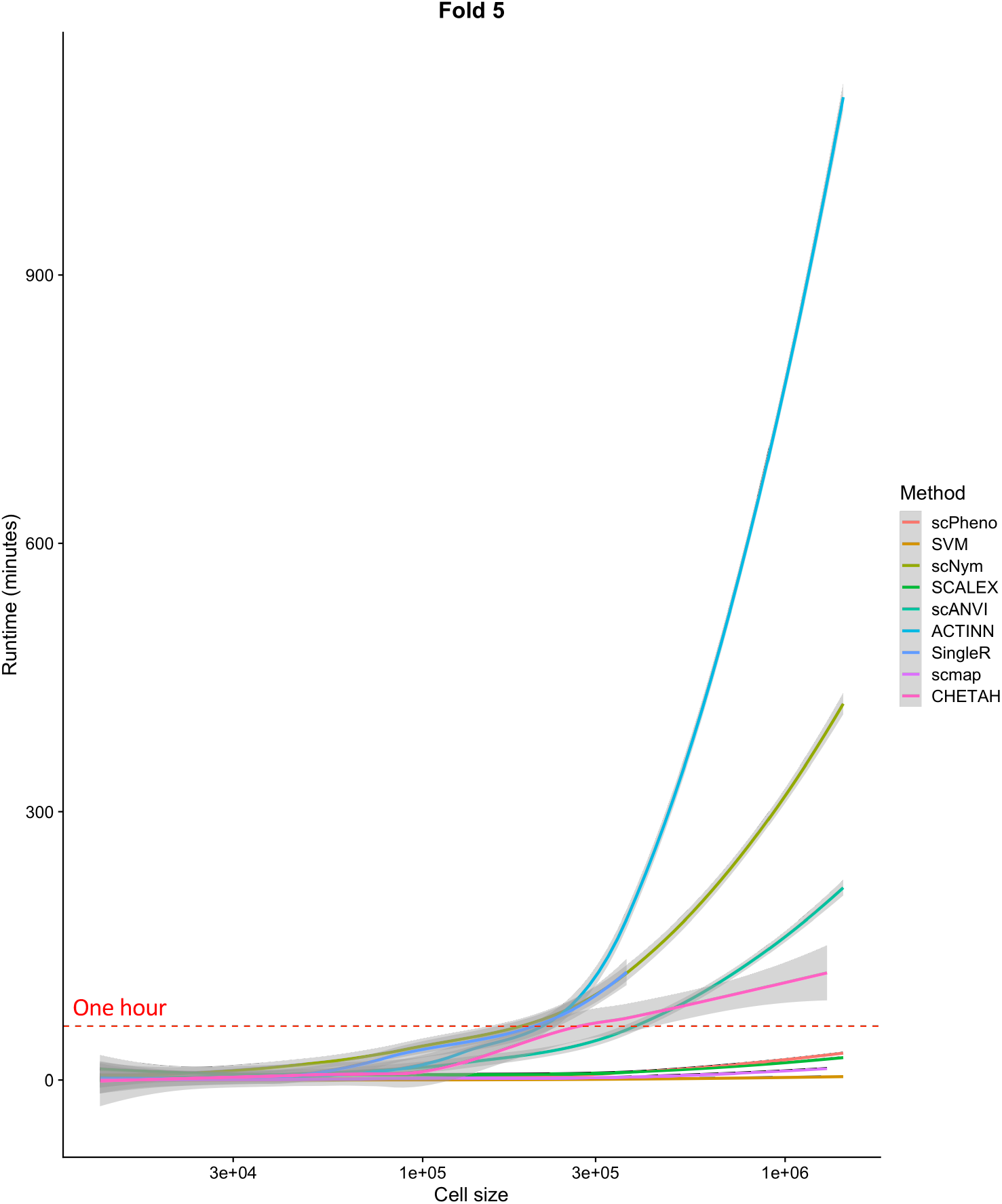
Runtime of the five-fold evaluation experiment including the training time and the validation time. The red dashed line represents one hour.

**Supplementary Fig. 6.**
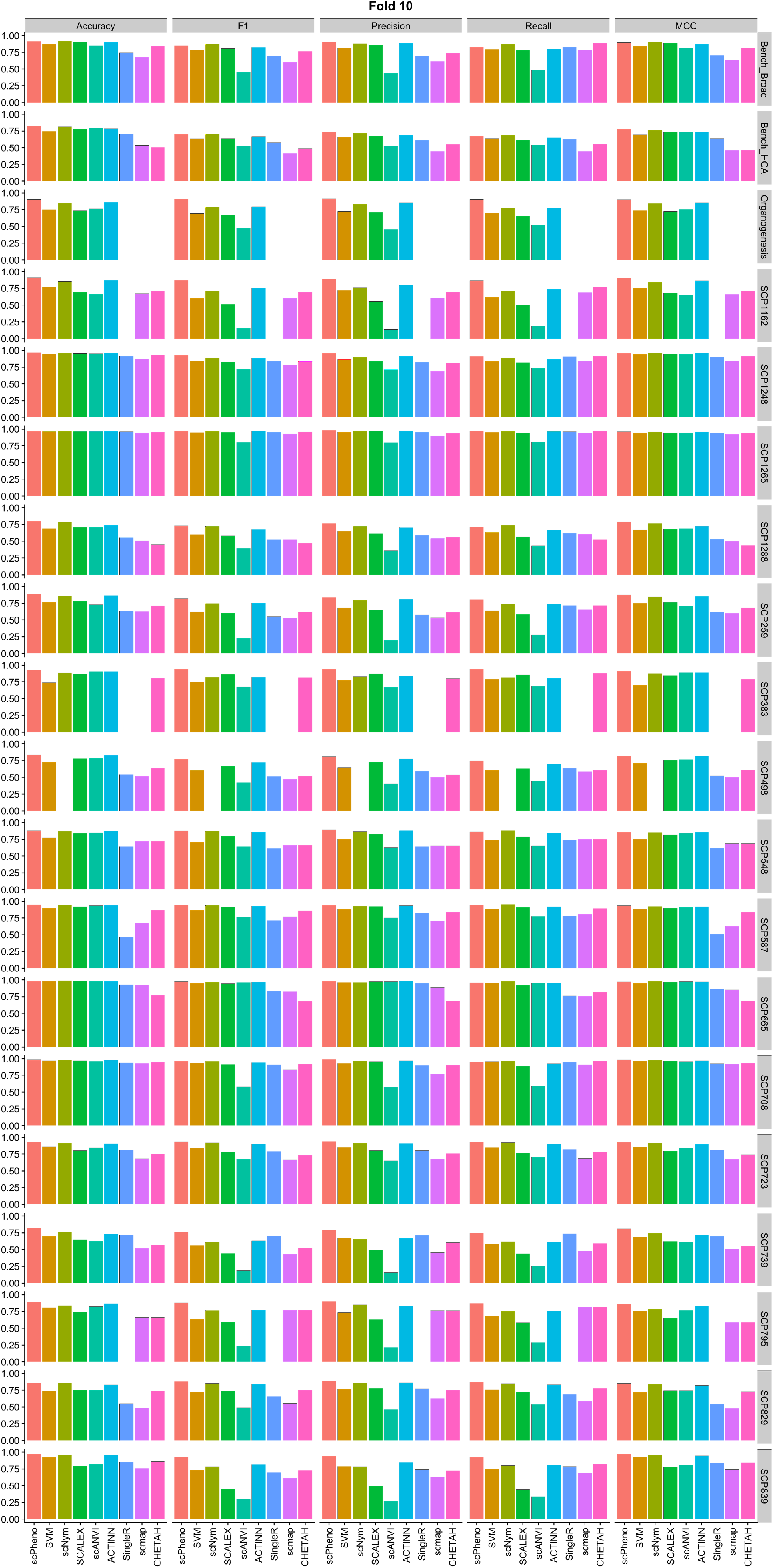
Evaluation of the cell-type prediction, ten-fold. For each benchmark dataset, 90 percent of the dataset were used for validation and the rest for training.

**Supplementary Fig. 7.**
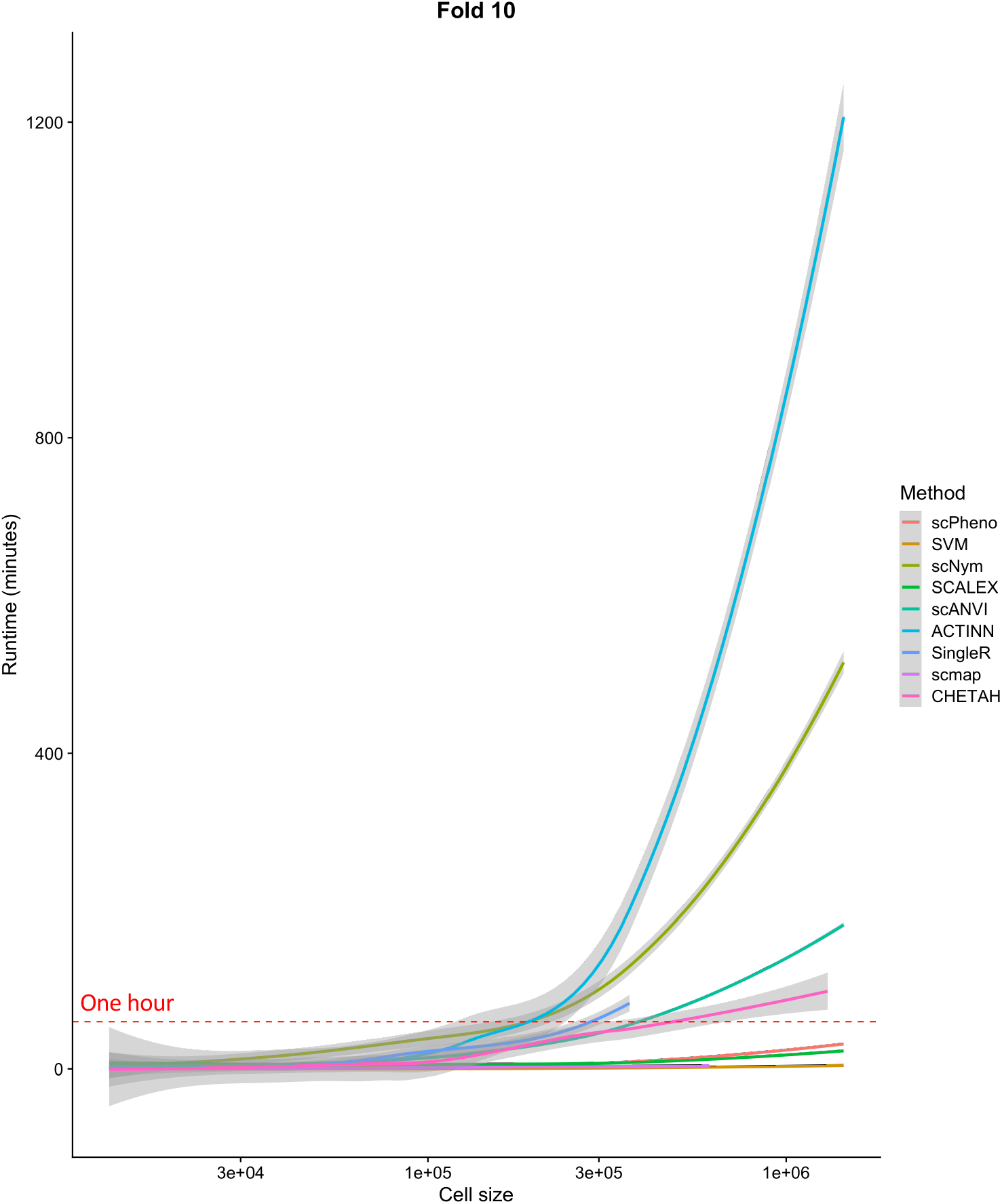
Runtime of the ten-fold evaluation experiment including the training time and the validation time. The red dashed line represents one hour.

**Supplementary Fig. 8.**
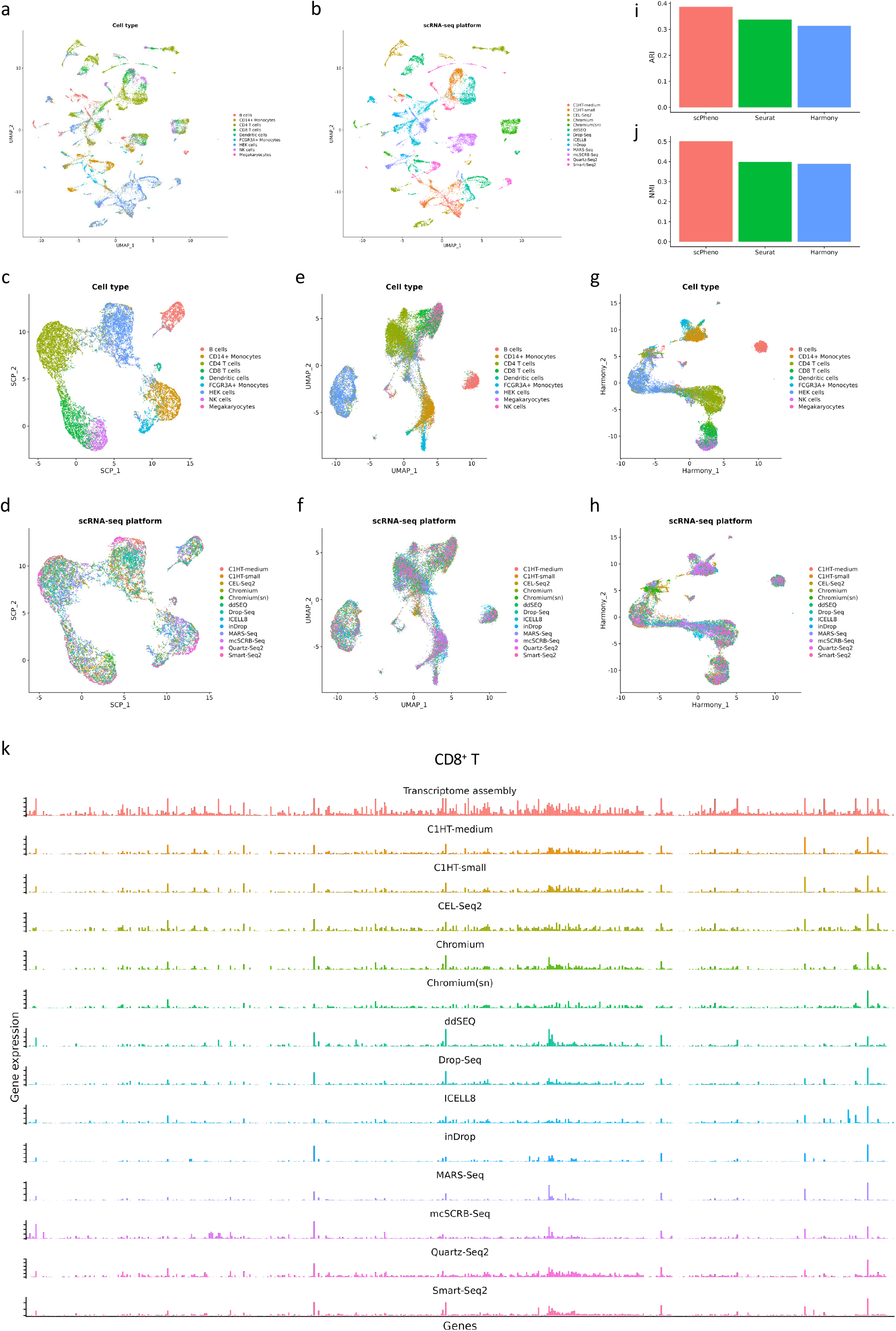
Evaluation of scRNA-seq integration with the Bench_HCA dataset. **a**, the visualization of cell types before integration. **b**, The visualization of scRNA-seq platforms before integration. **c**, The visualization of cell types after integration by scPheno. **d**, The visualization of scRNA-seq platforms after integration by scPheno. **e**, The visualization of cell types after integration by Seurat. **f**, The visualization of scRNA-seq platforms after integration by Seurat. **g**, The visualization of cell types after integration by Harmony. **h**, The visualization of scRNA-seq platforms after integration by Harmony. **i**, The adjusted Rand index (ARI) indicated how well cells of the same cell type were integrated. **j**, The normalized mutual information (NMI) evaluated the quality of the integration by removing the confounding factor of the size imbalance between cell types. **k**, Different scRNA-seq platforms display biases in digitalizing gene expressions on the transcriptome. scPheno can assemble the transcriptome through the estimation of gene expressions based on the integration of multiple scRNA-seq platforms (top panel).

**Supplementary Fig. 9.**
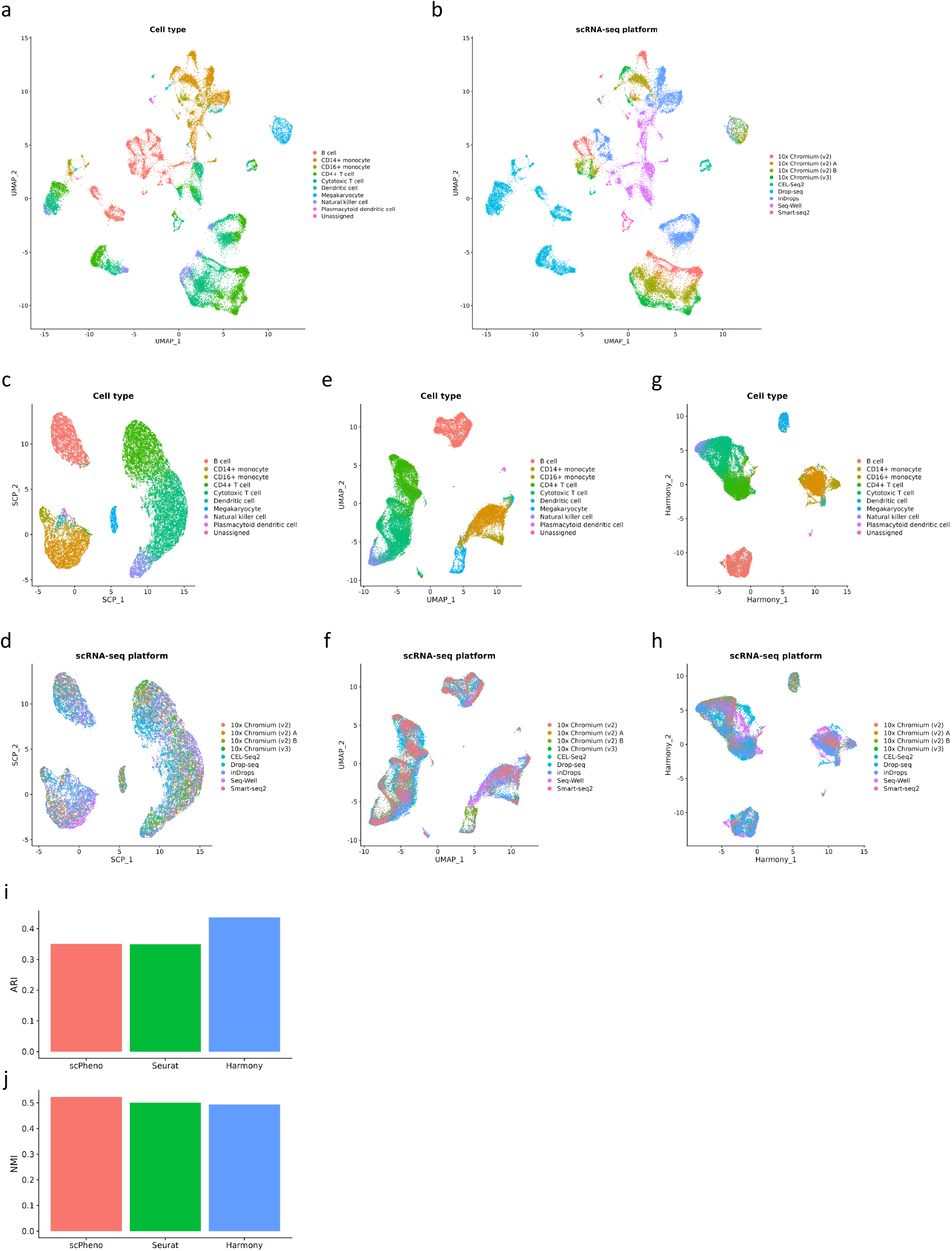
Evaluation of scRNA-seq integration with the Bench_Broad dataset. **a**, the visualization of cell types before integration. **b**, The visualization of scRNA-seq platforms before integration. **c**, The visualization of cell types after integration by scPheno. **d**, The visualization of scRNA-seq platforms after integration by scPheno. **e**, The visualization of cell types after integration by Seurat. **f**, The visualization of scRNA-seq platforms after integration by Seurat. **g**, The visualization of cell types after integration by Harmony. **h**, The visualization of scRNA-seq platforms after integration by Harmony. **i**, The adjusted Rand index (ARI) indicated how well cells of the same cell type were integrated. **j**, The normalized mutual information (NMI) evaluated the quality of the integration by removing the confounding factor of the size imbalance between cell types.

## References

1. Ma, Q. et al. Global Percentage of Asymptomatic SARS-CoV-2 Infections Among the Tested Population and Individuals With Confirmed COVID-19 Diagnosis: A Systematic Review and Meta-analysis. JAMA Netw. Open 4, e2137257–e2137257 (2021).

2. Wilk, A. J. et al. A single-cell atlas of the peripheral immune response in patients with severe COVID-19. Nat. Med. 26, 1070–1076 (2020).

3. Wang, G. et al. A deep-learning pipeline for the diagnosis and discrimination of viral, non-viral and COVID-19 pneumonia from chest X-ray images. Nat. Biomed. Eng. 5, 509–521 (2021).

4. Ahern, D. J. et al. A blood atlas of COVID-19 defines hallmarks of disease severity and specificity. Cell 185, 916–938.e58 (2022).

5. Ren, X. et al. COVID-19 immune features revealed by a large-scale single-cell transcriptome atlas. Cell 184, 1895–1913.e19 (2021).

6. Divij, M. et al. Deep immune profiling of COVID-19 patients reveals distinct immunotypes with therapeutic implications. Science (80-.). 369, eabc8511 (2020).

7. Schulte-Schrepping, J. et al. Severe COVID-19 Is Marked by a Dysregulated Myeloid Cell Compartment. Cell 182, 1419–1440.e23 (2020).

8. Lourda, M. et al. High-dimensional profiling reveals phenotypic heterogeneity and disease-specific alterations of granulocytes in COVID-19. Proc. Natl. Acad. Sci. 118, e2109123118 (2021).

9. Anthony, K. et al. Severely ill patients with COVID-19 display impaired exhaustion features in SARS-CoV-2–reactive CD8+ T cells. Sci. Immunol. 6, eabe4782 (2021).

10. Rha, M.-S. et al. PD-1-Expressing SARS-CoV-2-Specific CD8+ T Cells Are Not Exhausted, but Functional in Patients with COVID-19. Immunity 54, 44–52.e3 (2021).

11. Burkhardt, D. B. et al. Quantifying the effect of experimental perturbations at single-cell resolution. Nat. Biotechnol. 39, 619–629 (2021).

12. Dann, E., Henderson, N. C., Teichmann, S. A., Morgan, M. D. & Marioni, J. C. Differential abundance testing on single-cell data using k-nearest neighbor graphs. Nat. Biotechnol. (2021) doi:10.1038/s41587-021-01033-z.

13. Zhao, J. et al. Detection of differentially abundant cell subpopulations in scRNA-seq data. Proc. Natl. Acad. Sci. 118, e2100293118 (2021).

14. Kingma, D. P. & Welling, M. Auto-encoding variational bayes. 2nd Int. Conf. Learn. Represent. ICLR 2014 - Conf. Track Proc. 1–14 (2014).

15. Seok, L. J. et al. Immunophenotyping of COVID-19 and influenza highlights the role of type I interferons in development of severe COVID-19. Sci. Immunol. 5, eabd1554 (2020).

16. Hao, Y. et al. Integrated analysis of multimodal single-cell data. Cell 184, 3573–3587.e29 (2021).

17. Becht, E. et al. Dimensionality reduction for visualizing single-cell data using UMAP. Nat. Biotechnol. 37, 38–44 (2019).

18. Stephenson, E. et al. Single-cell multi-omics analysis of the immune response in COVID-19. Nat. Med. 27, 904–916 (2021).

19. Blei, D. M. & Jordan, M. I. Variational inference for Dirichlet process mixtures. Bayesian Anal. 1, 121–143 (2006).

20. Li, H. et al. Dysfunctional CD8 T Cells Form a Proliferative, Dynamically Regulated Compartment within Human Melanoma. Cell 176, 775–789.e18 (2019).

21. Yu, S., Yu, K., Tresp, V., Kriegel, H.-P. & Wu, M. Supervised Probabilistic Principal Component Analysis. in Proceedings of the 12th ACM SIGKDD International Conference on Knowledge Discovery and Data Mining 464–473 (Association for Computing Machinery, 2006). doi:10.1145/1150402.1150454.

22. Pedregosa, F. et al. Scikit-learn: Machine Learning in Python. J. Mach. Learn. Res. 12, 2825–2830 (2011).

